# Distinct Basket Nucleoporins roles in Nuclear Pore Function and Gene Expression: Tpr is an integral component of the TREX-2 mRNA export pathway

**DOI:** 10.1101/685263

**Authors:** Vasilisa Aksenova, Hang Noh Lee, Alexandra Smith, Shane Chen, Prasanna Bhat, James Iben, Carlos Echeverria, Beatriz Fontoura, Alexei Arnaoutov, Mary Dasso

## Abstract

Nuclear pore complexes (NPCs) are important for many processes beyond nucleocytoplasmic trafficking, including protein modification, chromatin remodeling, transcription, mRNA processing and mRNA export. The multi-faceted nature of NPCs and the slow turnover of their components has made it difficult to understand the role of basket nucleoporins (Nup153, Nup50 and Tpr) in these diverse processes. To address this question, we used an <u>A</u>uxin-<u>I</u>nduced <u>D</u>egron (AID) system to distinguish roles of basket nucleoporins: Loss of individual nucleoporins caused distinct alteration in patterns of nucleocytoplasmic trafficking and gene expression. Importantly, Tpr elimination caused rapid and pronounced changes in transcriptomic profiles within two hours of auxin addition. These changes were dissimilar to shifts observed after loss of Nup153 or Nup50, but closely related to changes after depletion of mRNA export receptor NXF1 or the GANP subunit of the TRanscription-EXport-2 (TREX-2) mRNA export complex. Moreover, GANP association to NPCs was specifically disrupted upon TPR depletion. Together, our findings demonstrate a unique and pivotal role of Tpr in regulating gene expression through GANP- and/or NXF1-dependent mRNA nuclear export.

## Introduction

Eukaryotic mRNA nuclear export requires a series of evolutionarily conserved complexes that bring nascent transcripts to the nuclear envelope and facilitate passage of messenger ribonucleoprotein (mRNP) particles through the nuclear pore complex (NPC). A key player in this process is the TRanscription and EXport 2 (TREX-2) complex, which bridges the transcription and export machineries in yeast through association to the Mediator complex, and the NPC^1^. TREX-2 associates with a structure of the NPC called the nuclear basket that protrudes from the nucleoplasmic face of the NPC^2,3^. The nuclear basket is comprised of three basket nucleoporins (BSK-NUPs) called Nup153, Tpr, and Nup50 in vertebrate cells^4^. The BSK-NUPs have been implicated in numerous processes beyond protein import and export, including chromatin remodeling, control of gene expression, and protein modification, as well as mRNA processing and export^5,6,7^. In particular, TREX-2 associates to NPCs in yeast through the Nup153 homologue, Nup1^3^. TREX-2 association with vertebrate NPCs requires Nup153 and Tpr but is independent of ongoing transcription^8,2^. The mechanisms of BSK-NUP-TREX-2 interactions and their consequences for gene expression remain poorly understood.

## Results and Discussion

It has been difficult to analyze discrete NPC functions in the absence of vertebrate BSK-NUPs; knockout of these genes is lethal for organisms, and their depletion by RNAi requires extended incubations, potentially allowing the emergence of secondary phenotypes from prolonged NPC disruption or defective post-mitotic NPC re-assembly^9–12^. To eliminate individual BSK-NUPs rapidly and selectively, we tagged Nup50, Nup153, and Tpr with an <u>A</u>uxin <u>I</u>nducible <u>D</u>egron (AID) and a <u>N</u>eon<u>G</u>reen (NG) fluorescent protein using CRISPR/Cas9 gene editing to biallelically introduce sequences encoding these tags at their endogenous loci in DLD1 cells (Fig. 1a, Supplementary Fig. 1a, b, d-o). To express the TIR1 protein, which drives degradation of AID-tagged proteins, we used CRISPR/Cas9 to biallelically insert sequences encoding <u>I</u>nfra-<u>R</u>ed Fluorescent <u>P</u>rotein (IFP) linked to Myc-tagged TIR1 through a cleavable P2A sequence at the C-terminus of the ubiquitously expressed RCC1 protein (Regulator of Chromosome Condensation 1) (Supplementary Fig. 1c). The resulting RCC1-IFP-TIR1 fusion protein was rapidly cleaved to yield active TIR1 and a RCC1-IFP fusion protein (RCC1^IFP^) that we employed as a marker for chromatin in live imaging experiments. We will call the resulting cell lines DLD1-Nup50^NG-AID^, DLD1-Nup153^NG-AID^, and DLD1-^AID-NG^Tpr in this report.

**Figure 1.**
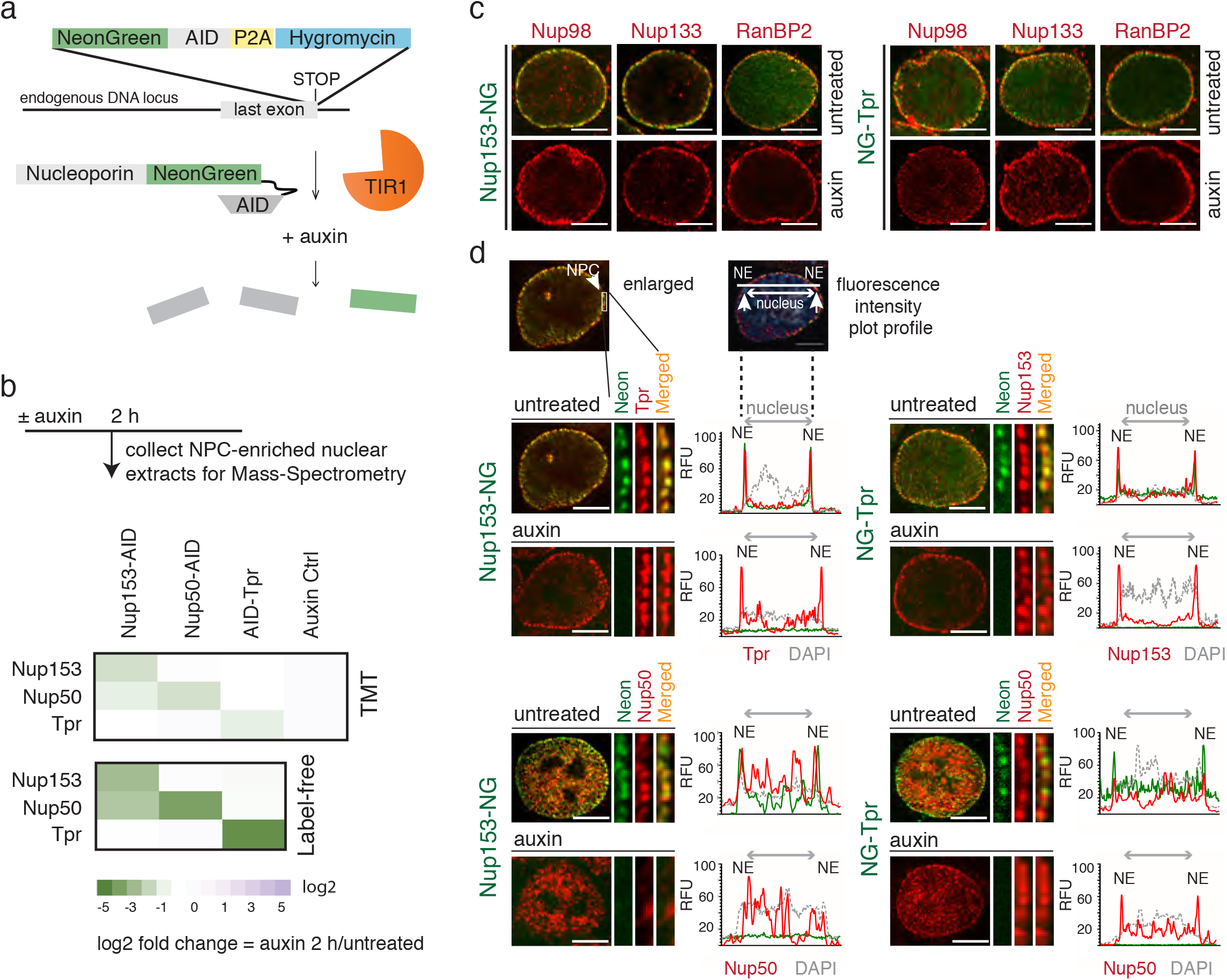
Stability of the assembled nuclear pore upon rapid loss of BSK-NUPs. **a**, A strategy of CRISPR/Cas9-based tagging of nucleoporin genes with AID sequence and degradation of the AID-fused proteins. **b**, Heat maps of differential abundance of Nup153, Nup50, and Tpr proteins in NPC-enriched nuclear extracts of indicated AID-tagged cells 2 h after auxin treatment using TMT-assisted and conventional (Label-free) mass-spectrometry. **c**, Localization of Nup98, Nup133, and RanBP2 in the absence or presence of Tpr and Nup153 (4 h of auxin treatment). **d**, Localization of BSK-NUPs in the absence or presence of Tpr and Nup153 (4 h of auxin treatment). Plots show line scans of relative fluorescence intensity (RFU) for BSK-NUPs and DNA across the nucleus. DNA was counterstained with DAPI. Note that Nup50 no longer localizes to the NE in the absence of Nup153. NE – nuclear envelope. Scale bar is 5 *μ*m.

Without auxin, DLD1-Nup50^NG-AID^, DLD1-Nup153^NG-AID^, and DLD1-^AID-NG^Tpr cells were viable and did not show any overt defects. We confirmed that all AID-tagged fusion proteins had the expected nuclear rim localization (Supplementary Fig. 2a), their expression did not alter the assembly of other NPC structures (Fig. 1c), and they became undetectable within one hour of auxin addition (Supplementary Fig. 1f, j, n). To assess how prolonged nucleoporin deficiency impacts cell survival^12–14^, we examined viability of DLD1-Nup50^NG-AID^, DLD1-Nup153^NG-AID^, and DLD1-^AID-NG^Tpr cells in the continuous presence of auxin (Supplementary Fig. 1g, k, o). Nup50-depleted cells continued to divide, albeit somewhat more slowly than control DLD1-Nup50^NG-AID^ cells. However, depletion of Nup153 or Tpr caused growth arrest, indicating that these two nucleoporins are essential for cell growth and proliferation, while the presence of Nup50 is less critical.

NPCs assemble as mammalian cells exit from mitosis and remain relatively stable through interphase until cells enter the next prophase^15, 16^. Due to the multiple cell cycles required to achieve BSK-NUP depletion in earlier experiments utilizing RNAi, changes in NPC composition observed in those experiments reflect interdependence of nucleoporins both for recruitment during post-mitotic NPC assembly and for persistence within assembled NPCs. The rapidity of AID-mediated degradation allowed us to assess the stability of nucleoporins within assembled NPCs. We assayed localization of GLFG nucleoporin Nup98, Y-complex component Nup133, and cytoplasmic fibril component RanBP2, which reside in different domains of the NPC. All of these nucleoporins maintained their localization after depletion of BSK-NUPs (Fig. 1c, Supplementary Fig.4a), and loss of individual BSK-NUPs did not cause visible nuclear envelope deformation. Our findings thus argue that loss of BSK-NUPs from assembled NPCs does not grossly impair nuclear pore architecture or composition in other NPC domains.

Earlier observations indicate that Tpr is dispensable for Nup153 and Nup50 localization^17,18^, and that Nup50 is likewise dispensable for Nup153 and Tpr localization^10,12^. However, there are conflicting reports regarding whether Nup153 depletion displaces Tpr from NPCs^9, 10, 11, 2, 19^. To address this question, we examined BSK-NUPs by tandem mass tag (TMT) and conventional MALDI-TOF mass spectrometry, immunostaining (Fig. 1b, d, Supplementary Fig. 4b, c), and live imaging of cells in which other nucleoporins were tagged with mCherry at their endogenous loci (Supplementary Fig. 3b-h). Under these conditions, the localization of BSK-NUPs were remarkably independent; none of them were redistributed upon the loss of others, with the exception of Nup50 dispersion into the nucleoplasm upon Nup153 loss (Fig. 1d (lower left), Supplementary Fig. 3a, h, Supplementary Fig.4d). Importantly, Tpr remained at NPCs even after several hours of Nup153 depletion, although we observed that post-mitotic recruitment of Tpr to NPCs was defective in the absence of Nup153 (Supplementary Fig.4e, f).

We assessed whether protein import and export were affected in cells lacking BSK-NUPs. We tested Importin-β dependent protein import^20^ and Crm1-dependent export^21^ using model substrates (Supplementary Fig. 5a, c). For comparison, we also examined similarly constructed cells with AID-tagged RanGAP1 (DLD1-RanGAP1^NG-AID^ cells; Supplementary Fig. 2c-f), the principal GTPase activating protein for Ran-dependent nuclear transport^22^. After auxin addition, DLD1-RanGAP1^NG-AID^ cells showed drastic inhibition of both import and export, as expected. Rates of import and export did not change in DLD1-Nup50^NG-AID^ cells after auxin addition, while DLD1-^AID-NG^Tpr cells showed slower export, and DLD1-Nup153^NG-AID^ showed slower trafficking in both directions (Supplementary Fig 5b, d). Interestingly, these results collectively indicate that BSK-NUPs perform distinct and non-equivalent functions in nuclear trafficking. Even with these changes, cells depleted of BSK-NUPs maintain much higher protein import-export levels within 2 hours of depletion than cells depleted of RanGAP1.

To assess the role of BSK-NUPs in gene expression, we compared RNA-seq profiles upon their loss (Fig. 2a, b) and found surprisingly distinct transcriptomic profiles for each nucleoporin. There were significant changes in the abundance of RNA from 167 genes upon Tpr loss (differential expression (DE) ≥ 30%, adj.p-value < 0.05), 142 of which were Tpr-specific. Nup153 and Nup50 loss caused DE of 28 (10 Nup153-specific gene) and 78 (45 Nup50-specific genes) genes, respectively, without extensive overlap between the DE profiles (Fig. 2b, c). To determine how blocking nuclear-cytoplasmic trafficking alters RNA abundance, we compared the impact of BSK-NUP loss to defects in auxin-treated DLD1-RanGAP1^NG-AID^ cells. Among 154 RNAs that changed response to RanGAP1 loss, 111 were specific to DLD1-RanGAP1^NG-AID^ cells. Remarkably, majority of BSK-NUPs-regulated RNAs did not overlap with the changes observed in RanGAP1-depleted cells, indicating that BSK-NUP-dependent RNAs are not differentially expressed because of a simple deficit in bulk protein nuclear trafficking.

**Figure 2.**
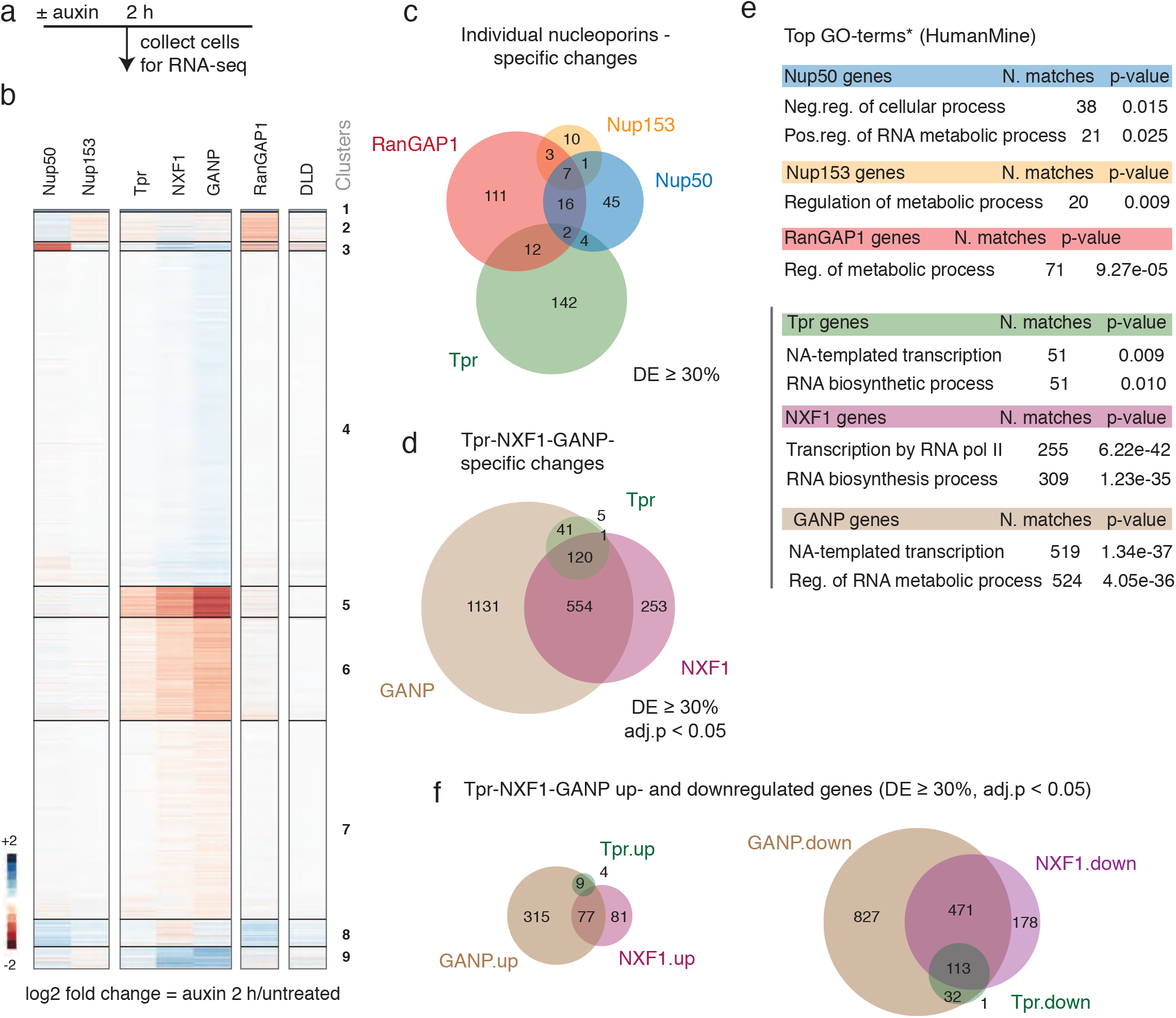
Loss of Tpr leads to rapid changes in mRNA abundance. **a, b**, A scheme of the experiment and heatmaps of unsupervised k-means clustering of differentially expressed genes 2 h after auxin treatment of cells expressing corresponding AID-tagged proteins. DLD – parental DLD1 cells treated with auxin. **c**, Venn diagram representing the number of RNAs that showed significant change (both up- and down-regulation) upon Nup50, Nup153, Tpr, and RanGAP1 loss. Some RNAs altered after loss of multiple nucleoporins are not shown to avoid confusion. The corresponding RNAs in each category: Nup153-Nup50-Tpr (1 gene), Nup50-Tpr-RanGAP1 (2 genes), Nup153-Tpr-RanGAP1 (1 gene), Nup153-Nup50-Tpr-RanGAP1 (2 genes). **d**, Venn diagram representing the number of significantly regulated transcripts upon Tpr, NXF1, or GANP depletion (p-value = 2.3e-117, Hypergeometric distribution test). **e**, Top GO-terms (*Biological Process) of individual differentially expressed (DE) RNAs upon loss of indicated nucleoporins, NXF1, or GANP (DE ≥ 30 %, adj. p-value < 0.05, Wald test). **f**, Venn diagram representing overlaps between significantly up- or down-regulated transcripts upon Tpr, NXF1, or GANP loss. Note that most of the Tpr/NXF1/GANP-dependent RNAs are downregulated upon Tpr, NXF1, or GANP loss.

Direct analysis of RNA abundance after Nup50 or Tpr depletion has not been documented. While altered transcriptomic profiles of mouse embryonic stem cells^11^ and Drosophila SL-2 cells ^23^ after Nup153 depletion using shRNAs have been reported, it is difficult to compare our data directly to those findings because of experimental differences, particularly the extended time required for shRNA-based depletion. Importantly, our data allow us to identify rapid, BSK-NUP-dependent transcriptomic changes that are unlikely to arise through altered post-mitotic NPC assembly defects or prolonged disruption of nuclear transport. We focused on Tpr-specific RNAs because Tpr loss caused the greatest impact on gene expression among BSK-NUPs. We grouped differentially expressed genes in visually distinct clusters (k-means clustering; Fig. 2b, Supplementary Table 2). Within the Tpr-regulated RNAs, the preponderance were downregulated. Gene Ontology (GO) analysis (Fig. 2e, Supplementary Table 3) revealed that Tpr loss differentially and specifically impacted RNAs involved in regulation of transcription; whereas RanGAP1, Nup153, and Nup50 specific-genes showed little or no enrichment in these categories (Fig. 2e). DLD1-^AID-NG^Tpr cells treated with Actinomycin D (ActD) did not show differential expression of Tpr-regulated RNAs in the presence versus absence of auxin (Fig. 4a, Supplementary Table 2), indicating that ongoing transcription was required for their relative change in abundance. At the same time, only 16 of the genes whose expression changed upon Tpr loss demonstrated Ser5P Pol II occupancy (p-value = 0.003) at the promoter or gene body in ChIP experiment (Fig. 4b, Supplementary Fig. 5e, Supplementary Table 4), arguing that transcription rates for most of these genes are not greatly altered after Tpr loss.

Importantly, long-term loss of Tpr led to measurable retention of bulk poly(A) mRNA in nuclear speckles that was not observed after Nup153 or Nup50 loss (Fig. 3a, Supplementary Fig. 5f-h), and mass spectrometry proteomic analysis of nuclear-pore associated proteins showed a reduction of GANP (germinal center-associated nuclear protein, a TREX-2 scaffolding subunit)^24^ in Tpr-depleted cells, but not Nup153- or Nup50-depleted cells (Supplementary Fig. 4h, Supplementary Table 1). Notably, GANP-depletion causes polyA RNA accumulation in nuclear speckles^25,2,26^, reminiscent of our observations in Tpr-depleted cells. Depletion of the mRNA export factor NXF-1 likewise causes polyA RNA accumulation in nuclear speckles^2^, and Tpr depletion disrupts both TREX-2^2,27^ and NXF-1-dependent RNA export^25^. To further examine the relationship between these RNA export factors and Tpr, we engineered AID-tagged cell lines for GANP (DLD1^AID-NG^GANP, Supplementary Fig. 2l-o), and NXF1 (DLD1^AID-FLAG^NXF1, Supplementary Fig. 2g-j).^AID-FLAG^NXF1 and ^AID-NG^GANP were degraded within one to three hours of auxin addition, resulting in a rapid arrest of cell growth and loss of cell viability (Supplementary Fig.2 h, m).

**Figure 3.**
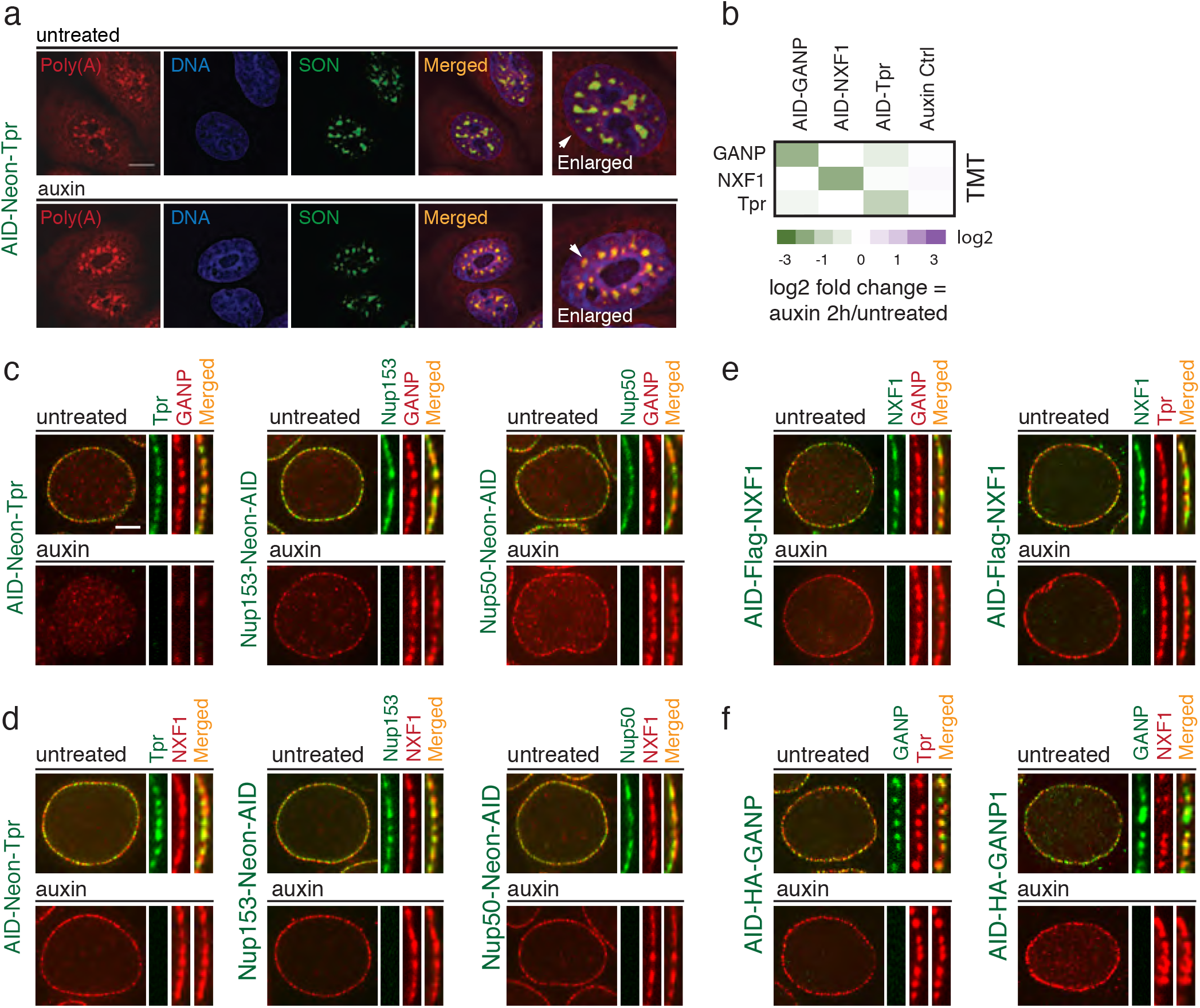
Tpr is required for GANP but not NXF1 tethering to the NE and poly(A) RNA nuclear export. **a**, The nuclear-cytoplasmic distribution of poly(A) RNA in AID-NGTpr targeted cells using oligo(dT)-Quasar 670 probe 8 h after Tpr degradation. Poly(A) RNA accumulates in the nuclear speckles of auxin-treated Tpr cells. Scale bar, 10 *μ*m. **b-c**, Tpr loss mislocalizes GANP from the NE. **b**, A heat map of TMT-based mass-spectrometry of NPC-enriched nuclear extracts shows changes in GANP, NXF1, and Tpr abundance 2 h after auxin treatment. c, GANP localization at the NE depends on Tpr, but not on Nup153 or Nup50. **d**, NXF1 localization is not affected upon Tpr, Nup153, or Nup50 loss. **e, f**, GANP and NXF1 localization at the NE are independent of each other. Correspondent AID-tagged cell lines were treated with auxin for 2 h. Scale bar is 5 *μ*m.

We compared the impact of TREX-2 or NXF1 depletion to the loss of BSK-NUPs, analyzing both protein localization and changes in the transcriptome. Immunofluorescent microscopy showed that GANP is dispersed after Tpr loss, but not after Nup50 or Nup153 depletion (Fig. 3c), confirming the changes indicated by mass spectrometry (Fig. 3b). GANP and NXF1 were each retained at the NPC in the absence of the other, and Tpr localization was not dependent on either protein (Fig. 3e-f). Transcriptomic analysis showed that Tpr-dependent RNAs overlapped almost entirely with GANP-dependent transcripts, which in turn overlapped extensively with NXF1-dependent RNAs (Fig. 2d, p-value = 2.3e-117) with similar patterns of up- and down-regulation (Fig. 2b, f). GANP, NXF1, and Tpr specific RNAs share similar GO terms (Fig. 2e, Supplementary Table 3). In combination with the largely dissimilar transcriptomic patterns observed after Nup153 or Nup50 depletion, these data suggest that Tpr has a unique function within the NPC basket to facilitate export of TREX-2- and NXF1-dependent RNAs.

We examined changes of individual Tpr-regulated RNAs in more detail. *fjx1* and *c-fos* transcripts were chosen as examples of down- and up-regulated mRNAs (Fig. 4c-e). We confirmed that *fjx1* mRNA decreased after Tpr depletion by qPCR (Fig. 4e) and analyzed Ser5P Pol II distribution by ChIP-Seq (Fig. 4c); *fjx1* showed lowered mRNA levels but little change in Ser5P Pol II ChIP-Seq patterns, suggesting that the changes in *fjx1* abundance occurred post-transcriptionally. To assess its retention in the nucleus, we performed *in situ* analysis using a hybridization-based signal amplification protocol that allows single molecule visualization with low background signal^28^: *fjx1* transcripts were less abundant after Tpr depletion, and retained within the nucleus with the ratio of cytoplasmic to nuclear signal (0.29±0.04, Fig. 5b) that was significantly different (p-value = 0.0003) from the level observed in untreated cells (1.3±0.08). Taken together, this pattern suggested that *fjx1* mRNA may be transcribed at a similar rate in the absence of Tpr, but that it is not productively exported, potentially leading to more rapid turnover. We similarly confirmed by qPCR that *c-fos* mRNA was up-regulated in response to Tpr depletion (Fig. 4e). In this case, Ser5P Pol II distribution by ChIP-Seq (Fig.4c), and *in situ* hybridization indicated an increase in ongoing transcription of *c-fos* (Fig. 5a). Interestingly, like *fjx1, c-fos* transcripts showed a strong decrease in the ratio of cytoplasmic to nuclear signal (0.39±0.09, Fig. 5b) in comparison (p-value = 0.0022) to untreated cells (1.4+0.17), again suggesting that they were not effectively exported. Notably, the distribution patterns of both mRNAs closely mimicked their patterns in the absence of GANP or NXF1 (Fig. 5a). We thus observed a consistent pattern in which both down- and up-regulated mRNAs fail to be translocated to the cytoplasm in the absence of Tpr, GANP or NXF1.

**Figure 4.**
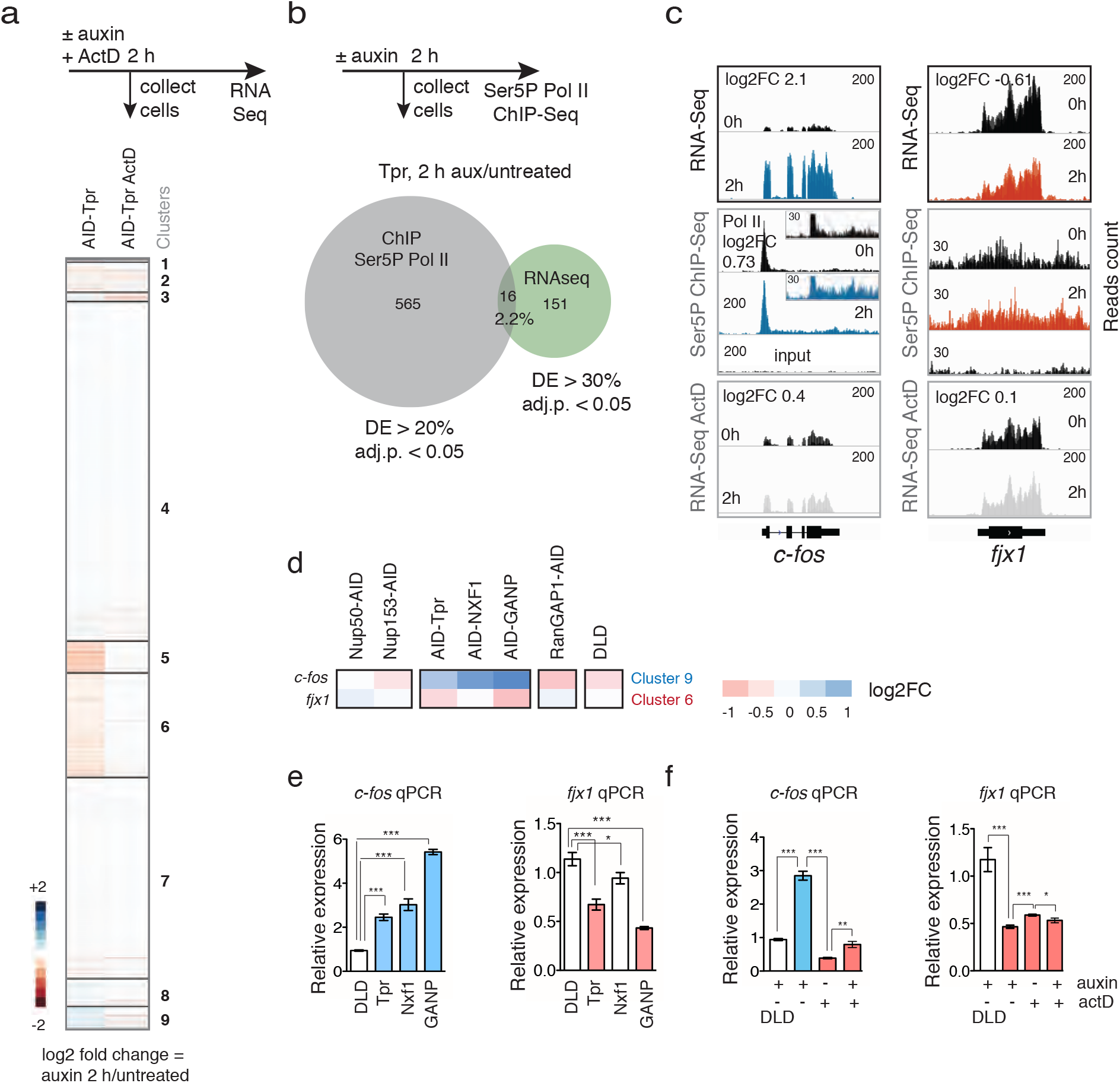
Tpr-regulated RNAs rely on ongoing transcription. **a**, Heat map of differential RNA expression upon Tpr loss with and without ongoing transcription: Left lane shows a heat map of differentially expressed RNAs upon loss of Tpr (2 h auxin vs. untreated). Right lane shows differentially expressed RNAs upon loss of Tpr in the presence of ActD (2 h ActD and auxin vs. ActD). **b**, A Venn diagram of Ser5P Pol II enriched genes and differentially expressed RNAs 2 h after Tpr loss. Pol II occupancy around the transcription start sites (TSSs) and gene body was changed only for 16 differentially expressed Tpr-specific genes. *c-fos* and *fjx1* are examples of genes that increased or showed no change in Pol II occupancy, respectively. **c**, IGV snapshots of RNAseq, Ser5P Pol II ChIP-Seq, and ActD RNA-Seq for *c-fos* and *fjx1* transcripts. **d-f**, RNAseq (**d**) and RT-qPCR (**e-f**) assays of change of *c-fos* and *fjx1* RNAs 2 h after Tpr or GANP loss. Error bars are SD. Asterisks indicate p-value * < 0.1, ** < 0.01, *** < 0.001 (unpaired t-test), ns – non-significant. FC – Fold Change.

**Figure 5.**
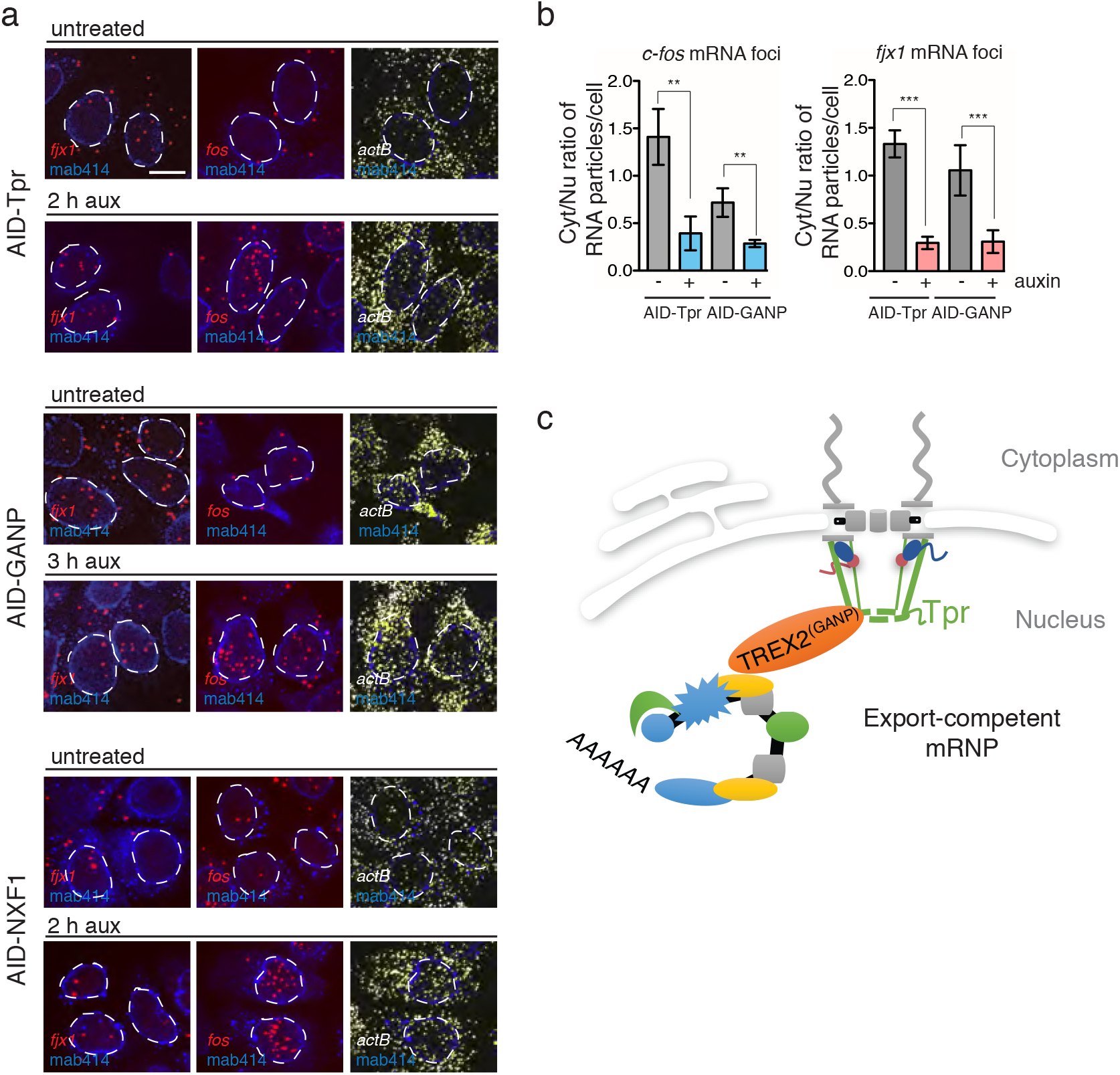
Tpr regulates RNA localization and abundance of GANP-dependent transcripts. **a, b**, *fjx1, c-fos*, and *actb* RNA localization 2 h after Tpr, GANP, or NXF1 loss (**a**). Ratio of cytoplasmic vs. nuclear (Cyt/Nu) *fjx1* and *c-fos* transcripts after Tpr or GANP loss (**b**). Asterisks indicate p-value ** < 0.01, *** < 0.001 (unpaired t-test). **c**, Schematic representation of Tpr’s role in gene expression through TREX-2-dependent mRNA export. Tpr anchors GANP to the NPC, and Tpr loss causes retention of GANP-dependent RNAs within the nucleus which affects their stability.

In summary, rapid depletion of AID-tagged BSK-NUPs allowed us to demonstrate discrete functions of BSK-NUPs in NPC organization, nuclear trafficking, and gene expression with a temporal resolution that was previously not attainable. This system will be extremely useful in future analysis of many other transport and signaling pathways that are associated to the NPC, not only in discovering which nucleoporins act in these contexts, but also in discovering how the NPC may act as a hub for integration of these pathways to each other. Notably, we found that BSK-NUPs are targeted to the nuclear pore independently of each other, except Nup50 localization which depends on Nup153. Loss of Nup50, Nup153, or Tpr led to unique transport and transcriptomic changes that were very different from protein import-export defects mediated by RanGAP1 loss. Remarkably, Tpr nucleoporin-specific changes in RNA abundance closely mimic NXF1 or GANP loss, indicating an integral role of Tpr in TREX-2- and NXF1-dependent RNA export (Fig.5c). Consistent with this idea, Tpr was the only anchoring nucleoporin within the basket required for recruitment of GANP subunit of TREX-2 complex to the NPC, further demonstrating an exclusive function of Tpr in coupling transcription and export that is not shared with Nup153 or Nup50. We speculate that the requirement for Nup153 in recruitment of Tpr during post-mitotic NPC re-assembly may at least partially explain previous observations arguing for an essential role of Nup153 in TREX-2-dependent RNA export^2^. Given the singular importance of Tpr in this context, it will be important to investigate in future experiments whether it simply acts as a landing platform for TREX-2 at the NPC or actively participates in remodeling or translocation of mRNPs in transit.

## Methods

### Cell culture and gene targeting

The human colorectal cancer cell line DLD-1 was cultured in DMEM (Life Technologies) supplemented with heat-inactivated 10% FBS (Atlanta Biologicals), antibiotics (100 IU/ml penicillin and 100 μg/ml streptomycin), and 2 mM GlutaMAX (Life Technologies) in 5% CO_2_ atmosphere at 37°C. For transfection 5*10^4^ cells/well were plated in 24-well plates a day before transfection. Plasmids for transfection were prepared using the NucleoSpin buffer set (Clontech) and VitaScientific columns. Plasmids were not linearized before transfection. Cells for gene targeting were transfected with 500 ng of donor and gRNA plasmids in ratio 1:1 using ViaFect (Promega) transfection reagent according to the manufacturer’s instruction. Seventy-two hours after transfection, cells were seeded on 10-cm dishes with the selective antibiotics (hygromycin 200 μg/ml, blasticidin 10 μg/ml or puromycin 3 μg/ml) until clones were formed on a plate. Analysis of localization and expression of targeted proteins in clones were performed after 14- to 20-day period post transfection on a 24-well lumox (Sarstedt) plates. Clones with proper protein localization were propagated for genomic PCR and Western blot analysis in regular complete media without selective antibiotics.

Regulator of chromosome condensation 1 (RCC1, NC_000001.11) locus was chosen to knock-in TIR1. After CRISPR/Cas9-mediated recombination, RCC1 was targeted with an infra-red fluorescent protein (IFP2.0 Addgene #54785), TIR1 was targeted with 9 Myc-tag sequences, and both proteins were separated from each other and from a protein encoded by blasticidin resistance gene (cloned from pQCXIB #631516, Clontech) by self-cleavage peptide P2A. Only bright, healthy growing clones were propagated for downstream analysis. Because TIR1 integration potentially can reduce protein level of AID-targeted protein we first integrated AID degron and a NeonGreen fluorescent protein up- or downstream of basket nucleoporins’ genes, and then TIR1 in the rcc1 locus.

### Plasmid construction

The CRISPR/Cas9 system was used for endogenous genes targeting. All gRNA plasmids were generated with primers listed in Supplementary Table 5 (IDT) and integrated into pX330 (Addgene #42230) vector using Zhang Lab General Cloning Protocol ^29^. For gRNA selection we used CRISPR Design Tools from http://crispr.mit.edu:8079 and https://figshare.com/articles/CRISPR_Design_Tool/1117899. pEGFP-N1 vector (Clontech) was used to create a universal donor vector (pCassette). Briefly, the pEGFP-N1 vector was restricted with Nde1 and Not1 enzymes (NEB), and a new multiple cloning site was inserted into the vector instead of sequences corresponding to the CMV promoter and EGFP protein. The sequences of homology arms were amplified from genomic DNA extracted from DLD-1 cells, the full-length 229-amino acid AID degron (flAID), hygromycin, and TIR1 sequence were amplified by PCR from pcDNA5-EGFP-AID-BubR1 (Addgene #47330) and pBABE TIR1-9Myc (Addgene #47328)^30^ plasmids. The DNA sequence of NeonGreen fluorescent protein, FLAG tag, HA tag, a minimal functional AID tag (1xmicroAID) 71-114 amino-acid^31^, three copies of reduced AID tag (3xminiAID) 65-132 amino-acid^32^ were codon optimized and synthesized in IDT company. Tpr, Nup153, Nup50, NXF1 were tagged with flAID, RanGAP1 with 3xminiAID and GANP with 1xmicroAID tags (Supplementary Fig. 1, 3). cDNAs of mCherry and puromycin resistance were amplified from pmCherry-N1 (632523, Clontech) and pICE vector (Addgene #46960)^33^, respectively. All PCR reactions were performed using Hi-Fi Taq (Invitrogen) or Herculase II Fusion (Agilent) DNA polymerases with primers listed in Supplementary Table 5.

### Genotyping

DNA from DLD-1 and nucleoporins-targeted cells was extracted with Wizard^®^ Genomic DNA Purification Kit (Promega). Clones were genotyped by PCR for homozygous insertion of tags with two sets of primers listed in Supplementary Table 5.

### Crystal violet staining

Survival of AID-targeted nucleoporins after auxin treatment was estimated with crystal violet staining. The procedure was based on Feoktistova^34^ with modifications. In brief, suspension of 4-5*10^3^ cells/well was plated in twelve independent wells of 96-well plate. After cells were attached to the plastic, fresh media with 1 mM auxin was added into the corresponding wells, followed by incubations for 0 h, 24 h, 48 h, and 72 h. Zero-hour time point was used for cell number normalization between different cell lines. Cells were washed with PBS, fixed with 70% ethanol for 15 min, stained with 0.1% crystal violet for 15 min, and washed four times in a stream of distilled water. A hundred microliters of 10% acetic acid were added to each well with cells and incubated for 15 min at RT. Measurement of optical density was performed on EnSpire Plate Reader (Perkin Elmer) at 570 nm (OD570). To calculate the number of survived cells, we plotted the calibration curve with the estimated number of cells and measured their OD570.

### Time-lapse fluorescence microscopy

DLD-1 cells were grown on 4-well glass bottom chambers (Ibidi), imaged on the Eclipse Ti2 inverted microscope (Nikon), equipped with an Ultraview spinning disk confocal system (Ultraview Vox Rapid Confocal Imager; PerkinElmer) and controlled by Volocity software (PerkinElmer) utilizing Nikon CFI60 Plan Apochromat Lambda 60x/1.4 oil immersion objective lens with D-C DIC slider 60x II, and 40x/1.3 oil Nikon PlanFluor immersion objective lens. Cells were imaged once every 10 min in FluoroBrite DMEM (ThermoFisher) media. The microscope was equipped with temperature-, CO_2_- and humidity-controlled chamber that maintained a 5% CO_2_ atmosphere and 37°C.

NeonGreen and mCherry fluorescent protein siRNA accumulates in the nuclear specklesgnals were excited with a 488-nm (no more than 20% of power was applied) and 568-nm (no more than 50% of power was applied) laser lines, respectively. A series of 0.5 *μ*m optical sections were acquired every 10 min. Images were captured and analyzed using Volocity (PerkinElmer) and Image J (National Institutes of Health) software, respectively. Images represent maximum intensity projections of entire z-stacks.

### Immunofluorescence staining

DLD-1 cells expressing fluorescently targeted nucleoporins were seeded on coverslips in 12*10^4^ density on 60-mm dishes and grown for 2 days. Cells were washed with PBS, pH 7.4, and immediately fixed with 4% paraformaldehyde in PBS at RT for 15 min, permeabilized with 0.5% Triton X-100 for 10 min, and blocked with 10% horse serum for 20 min. For Nup133 antigen retrieval cells were washed with PBS and fixed with 4% paraformaldehyde in PBS with 0.5% Triton X-100 at RT for 15 min. For NXF1 and GANP nuclear envelope visualization cells were seeded on 8-well Nunc Lab-Tek slides and two days after washed three times with Buffer A (20 mM HEPES, pH 7.8; 2 mM DTT, 10% sucrose, 5 mM MgCl2, 5 mM EGTA, 1% TX100, 0.075% SDS) with the following 4% paraformaldehyde in PBS fixation at RT for 10 min. Localization of nucleoporins was detected by specific primary antibodies and AlexaFluor-conjugated secondary antibodies (Invitrogen). The incubation time for primary and secondary antibodies was 1 h. The nuclei were visualized with Hoechst (Invitrogen). Images were acquired using an Olympus IX71 inverted microscope (Olympus America Incorporation), equipped with an Ultraview spinning disk confocal system (Ultraview ERS Rapid Confocal Imager; PerkinElmer) and controlled by Volocity software (PerkinElmer) utilizing an Olympus UPlanSApo 100x/1.4 oil objective.

Brightness and contrast were applied equally to all images within the experiment using Fiji software version 2.0.0-rc-68/1.52e. RGB images from Fiji were processed to final resolution using Adobe Photoshop and Adobe Illustrator CS5.1.

### Analysis of NPC and NPC-associated proteins by mass spectrometry

Nuclear pore complexes-containing fraction (NPCCF) was isolated according to Cronshaw^35^ with modifications. Cells were seeded onto 150 mm plates and processed once they reach 90% confluency. Nocodazole (5 μg/ml) and Cytochalasin B (10 μg/ml) were added for 30 min and cells were washed twice with PBS. Cells were then incubated 2 x 5 min in 30 ml buffer A (20 mM HEPES, pH 7.8; 2 mM DTT, 10% sucrose, 5 mM MgCl2, 5 mM EGTA, 1% TX100, 0.075% SDS) at RT, followed by incubation in 15 ml buffer B (20 mM HEPES, pH 7.8; 2 mM DTT, 10% sucrose, 0.1 mM MgCl^2^), containing 4 μg/ml RNase A (Promega) for 10 min at 37°C, and washed in buffer A. NPCs and associated proteins were eluted by incubating the plates in 12 ml of buffer C (20 mM HEPES, pH 7.8; 150 mM NaCl, 2 mM DTT, 10% sucrose, 0.3% Empigen BB) for 10 min at 37°C. The supernatant was transferred to 14 ml polypropylene tubes and span for 3 min at 28 000g (HB-6 rotor) at 4°C. The supernatant was transferred to the new tubes, incubated on ice for 30 min, and saturated TCA was added to the final concentration of 8%. Tubes were vigorously mixed and incubated on ice for another 30 min. Precipitated protein complexes were sedimented by centrifugation at 28 000 g (HB-6 rotor) for 20 min at 4°C. The supernatant, containing some Empigen-TCA floating complexes, was aspirated. The protein pellet was resuspended in 1.5 ml cold (−20°C) ethanol, vigorously vortexed to solubilize Empigen-TCA complexes, transferred to eppendorf tubes and spun for 15 min at 11 000 g at 4°C using bucket rotor. The pellet was resuspended in ethanol and spun again for 1 min at 11 000 g. This wash was repeated twice. The NPCCF pellet was then solubilized in 100 μl of buffer D (8 M urea, 5 mM DTT) by pipetting, rigorously vortexed and centrifuged for 1 min at 11 000 g. The supernatant was collected, and the pellet was again solubilized in 100 μl of buffer D, vortexed and spun as above. Both supernatants were combined (200 μl total), and the protein concentration was measured. Samples were stored at −70°C before processing for mass spectrometry or SDS-PAGE.

For mass spectrometry analysis, 100 μg of NPCCF was resuspended in 50 μl of 8 M urea, supplemented with freshly added DTT (20 mM final) and incubated for 1 h at 37°C. Iodoacetamide (freshly made stock solution in 25 mM ammonium bicarbonate) was then added (50 mM final) and alkylation proceeded for 1 h at RT. The reaction was quenched by 50 mM DTT and samples were diluted 10 x with 25 mM ammonium bicarbonate. Three microgram of trypsin (v5111, Promega) was added and digestion reaction proceeded overnight at 37°C. Samples were acidified by formic acid, and peptides were desalted with Waters Oasis HLB 1cc columns. Peptides were eluted with 1 ml of buffer E (0.1% formic acid in 50% acetonitrile) and dried using SpeedVac. Samples were either individually analyzed with LCMS for label-free quantitation or analyzed as multiplex sets, after labeling with TMT reagents (TMT10plex label reagent set, Thermo Scientific), according to manufacturer’s instructions. To increase the protein coverage, each set of pooled TMT samples was separated into 24 fractions using basic reverse phase liquid chromatography (bRPLC)^36^.The mass spectrometric analysis was conducted on an LTQ Orbitrap Lumos (Thermo Fisher Scientific) based nanoLCMS system. Briefly, the label-free samples were analyzed with a 3 h gradient at a 120K resolution for MS1 scans and 30K resolution for MS2 scans at 30% HCD energy. Each TMT fraction was analyzed with a 2 h gradient at 120k resolution for MS1 and 50K for MS2 at 38% HCD energy.

Protein identification and quantitation analysis were carried out on Proteome Discoverer 2.2 platform (Thermo Fisher Scientific). Peptide IDs were assigned by searching the resulting LCMS raw data against UniProt/SwissProt Human database using the Mascot algorithm (V2.6, Matrix Science Inc.). And peptide-spectrum matches (PSM) were further validated with Percolate algorithm. Peptides with high confidence (<1% FDR) were filtered for protein identification. Label-free quantitation was based on areas under peptide precursor traces (MS1) that were aligned with the Minora feature detection function in PD2.2. For TMT sets, report ion intensities of ten channels were normalized at the total peptide amount level. Then relative quantitation of individual proteins was calculated using the normalized report ion intensities of unique peptides with <1% FDR and >30% isolation purity.

We determined the protein-level fold changes based on the median of peptide-level fold changes from the Proteome Discoverer-produced abundances in handling both TMT and Label-free results. Peptides that could be mapped to multiple proteins by Mascot were removed. We also discarded all keratin-related peptides based on the UniProt annotation. We separated peptides that mapped onto UniProt ID P52948 into NUP96 and NUP98, according to their mapping locations. Peptides that are mapped to amino acids from 1 to 880 were counted for NUP96; the others were used for NUP98. To minimize the batch effect, we used the quartile normalization before calculation of fold changes in the Label-free quantification. Such fold changes were visualized after k-means clustering. In determining the optimal k parameter for clustering, we used “mclust” package to calculate Bayesian Information Criterion of the expectation-maximization model^37^. The quartile normalization, k means clustering, and data visualization were performed under R development environment (R Core Team, 2018).

### Protein import and export assays

DLD-1 and nucleoporin-targeted cell lines were seeded on 8-well Nunc Lab-Tek slides, transfected with 500 ng of PXT_REV-GR-Turquoise2 (modified from Love et al., 1998)^20^, EGFP protein was replaced with Turquoise2, Addgene #60494^38^, or mCherry-LOV2-BiLINuS2^21^ plasmids using ViaFect (Promega) transfection reagent according to the manufacturer’s protocol. REV-GR-Turquiose2 and mCherry-LOV2-BiLINuS2 positive cells were imaged one to two days after transfection on Olympus IX71 inverted microscope as described in fluorescent microscopy section.

REV-GR-Turquoise2-transfected cells were treated with 100 nM Leptomycin B (LMB, LC Laboratories) for 30 min and then with 1 *μ*m Dexamethasone (Sigma Aldrich) for 10 min. The Turquoise2 protein was excited with a 405-nm laser line (50% of power). An exposure time of 500 ms was used to take images every 1 min for 10 min, as the import is complete within that time frame. The z-stack depth was roughly 8-12 *μ*m with a 1 *μ*m step size depending on the thickness of the cell.

The plates with mCherry-LOV2-BiLINuS2-transfected cells were covered by the aluminum foil for several hours before the assay. We noticed that any exposure to light before the initial background measurement negatively affected the data calculations. An image for background measurement, using 568 nm laser-line, at 20% power, was taken each time before blue light exposure. Adhesive 420 nm LED strips (1.6W, 100 Lumens, 21 LED LLC), attached to a large plastic disc (150 mm culture dish) were used to expose cells for 45 min and accumulate mCherry-LOV2-BiLINuS2 protein inside the nucleus. After blue light exposure, the light source was turned off, and one focal plane was imaged every minute for 10 min with the exposure time 500 ms.

Image analysis of nuclear import and export was performed on ImageJ software using Time Series Analyzer V3 plugin and ROI Manager dialog box.

For nuclear import measurements, we calculated an average from three points inside the cytoplasm and three points in nucleoli of each cell. The background value was subtracted from average values for the cytoplasm and the nucleus. The initial level of REV-GR-Turquiose2 protein in the nucleus was taken as a background for all downstream calculations. To calculate relative fluorescence intensity, we summarized the adjusted nuclear and cytoplasmic means, and then calculated the percentage of the total of the nuclear and cytoplasmic values using formula: 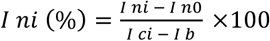, where *I_ni_* is intensity of the Turquiose2 protein in the nucleus of cell *i* after dexamethasone treatment multiplied by 100; *I_ni_* - initial level of Turquiose2 protein in the nucleus; *I_no_* - level of Turquiose2 proteinin the nucleus at time 0; *I_ci_* - total intensity level of Turquiose2 protein in the cell; *I_b_* - a background level of Turquiose2 protein in the cell *i*.

For nuclear export measurement, three points inside the nucleus of each cell were taken to calculate the mean of the nuclear signal at each time point. The initial level of mCherry-LOV2-BiLINuS2 inside the nucleus 45 min after blue light exposure was taken as 100% for all downstream calculations. The relative fluorescence intensity was calculated using formula: 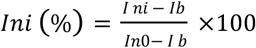, where *I_ni_* is intensity of the mCherry protein in the nucleus of cell *i* after blue light was turned off (multiplied by 100); *I_ni_* - initial level of mCherry protein in the nucleus; *I_no_* - level of mCherry protein in the nucleus at time 0; *I_b_* - a background level of mCherry protein in the cell *i*.

Variability in the data was calculated using the mean absolute deviation, which was calculated by taking the average of the absolute value of the difference between each data point and the corresponding median.

### Protein extraction and Western blot

Pellets of DLD-1 cells were lysed in 2x Laemmli Sample Buffer (Bio-Rad), boiled for 15 min at 98^0^C, and ultra-centrifuged at 117K for 10 min at 16^0^C. SDS-PAGE and Western blotting were performed as described elsewhere^39^. In brief, the protein samples were separated on 4-20% SDS-PAGE or Bolt™ 8% Bis-Tris gels (Invitrogen) for 1 h and blotted onto PVDF membrane. The membrane was blocked in 5% non-fat milk for 1 h with the following incubation with the primary antibody overnight at 4^0^C. Then the membrane was rinsed and probed for 2 h with the secondary anti-mouse or anti-rabbit antibodies conjugated to HRP with dilution 1:15000 in 1x TN buffer (150 mM NaCl, 10 mM TrisHCl, pH 7.5, 0. 1% Tween 20). Detection of the signal was performed using FluorChem Imaging System (ProteinSimple) with ECL Prime Western Blotting substrate (GE Healthcare) or SuperSignal WestPico PLUS Chemiluminescent Substrate (ThermoScientific).

### Quantitative PCR with reverse transcription (qRT-PCR)

Trizol-extracted RNA from cells was treated with Turbo DNA-free DNAse (ThermoFisher) and purified on RNAeasy MiniKit (Qiagen) columns according to the manufacturer’s protocol. One thousand three hundred micrograms of DNA-free RNA were used for cDNA synthesis with random hexamers (NEB) using SuperScript II (ThermoFisher) kit as suggested by the protocol. Quantitative PCR was performed using PrimeTime^®^ Predesigned qPCR Assays (IDT) Hs.PT.39a.22214847-ACTB, Hs.PT.39a.19639531-POLR2A, Hs.PT.58.22565110-HOXA13, Hs.PT.58.15540029-FOS, Hs.PT.58.40805543.g-EGR1, Hs.PT.58.40089589-GDF15, Hs.PT.58.19880529-ZNF576, Hs.PT.58.22812669-CLCF1, Hs.PT.58.194762.g-GADD45b, Hs.PT.58.40600236-TNFSF9, Hs.PT.58.2763744-METTL18, Hs.PT.58.22562604.g-OVOL1, Hs.PT.58.26129234.g-HIST1H2AB using CFX Connect™ Real-Time PCR Detection System (Bio-Rad). The relative transcript level was determined by normalizing to the expression level of ACTB and POLR2A genes using ddCq-method.

### RNA-Seq samples preparation

AID-targeted cells on 3^rd^-6^th^ passage were seeded on 6-well dishes for RNA isolation. Eight to ten hundred adherent DLD-1 cells were washed with PBS and lysed directly with RLT buffer from RNAeasy Mini Kit (Qiagen). Extracted RNA was analyzed for 260/280 and RIN value on Bioanalyzer 2100 (Agilent), only samples with RIN value more than 9 were proceeded for the library construction using Illumina TruSeq Stranded Total RNA Library Preparation Kit (Illumina) according to manufacturer’s instructions. All RNA samples were depleted for ribosomal RNA before library construction using Ribo-Zero™ Gold Kit H/M/R Kit (Illumina). Hi-Seq run of three independent biological replicates was performed on Illumina HiSeq2500. Twenty-thirty millions of 101 bp paired-end reads were generated per each replicate.

### RNA-Seq Analysis

We aligned the short-reads to the reference human genome GRCh38.p7/hg38 that we obtained from Ensembl (PMID:27899575). We used STAR aligner version 2.5.2b (PMID:23104886) for the mapping with default parameters. We quantified gene expression levels using feature Counts 1.4.6 (PMID:24227677). To obtain gene-level read counts, we supplied a gene annotation model from Ensemble Release 87, which corresponds to GENCODE version 25. We calculated gene-level FPKM values based on the longest-possible exon lengths after collapsing overlapping exons. For differential expression analysis, we used DESeq2 DESeq2 1.18.1 (PMID:25516281) as in the reference with one minor change; we applied expression cutoffs FPKM > 1 and Counts Per Million (CPM) > 1 before the analysis. We studied and visualized genes whose expressions are above the cutoff from more than 5% of our samples excluding for the Actinomycin D-treated samples. We used k-means clustering function in R environment 3.3.2 (R Code Team, 2018) to generate heat maps in the result section.

The Venn diagrams were created using Venny 2.1 tool and BioVinci software version 1.1.5, r20181005. Gene Ontology (GO-terms) enrichment analysis was performed using an integrated HumanMine database V5.1 2018 with Holm-Bonferroni test correction (p-value 0.05) for Biological Processes.

### ChIP-Seq Analysis

Chromatin immunoprecipitation assay with sequencing (ChIP-Seq) was performed using modified ChIP and ChIP-Seq protocols previously described ^40^, ^41^. Briefly, 3.5*10^6^ cells per 100 mm dish were washed with 1x PBS and cross-linked with 1% formaldehyde for 11 min. After quenching with 0.125 M glycine for 13 min, cells were scraped, lysed, and sheared by sonication using Bioruptor (Diagenode) for 42 min with 30 s pulse/pause cycles on ice. After enrichment of chromatin fragments around 300-bp, cell debris was spun down, and supernatant was incubated overnight at +4^0^C with anti-mouse IgG M-280 (ThermoFisher) dynabeads previously incubated with anti-Ser5P Pol II mouse antibodies (ab5408) for 10 h. Immune complexes were washed twice with IP, IP-500, LiCl, TE buffers and eluted using CHIP Elute Kit (Clontech). Two independent biological replicates were used for library construction using DNA SMART ChIP-Seq Kit according to the manufacturer’s protocol. In total 16 PCR cycles were used during library construction. About two hundred thousand million 50 bp paired-end reads were generated per each replicate. Alignment of short-reads was performed with BWA (VN:0.7.17-r1188) against reference human genome GRCh38.p7/hg38. Read quantitation over genes was tabulated using HOMER (PMID: 20513432). Differential IP recovery of genes was tested using DESeq2 (PMID: 25516281) and MACS2 broad peak calling tools. Plots were generated with IGV_2.4.6 software.

### RNA Fluorescent in Situ Hybridization (FISH)

poly(A)RNA FISH was performed as described earlier^42^. In brief, cells were treated with 1 mM auxin for 8 h, washed with PBS, fixed for 15 min in 4% PFA at room temperature (RT), incubated in 70% ethanol for 16h at 4 °C, washed with PBS, and incubated with wash buffer for 5 min at RT. Next cells were incubated with oligo^43^-Quasar 670 probes (LGC Biosearch Technologies) for 4h at 37 °C. Again cells were incubated with wash buffer at 37 °C for 30 min and washed with PBS. After that cells were stained with 1.5 μg/ml Hoechst 33258 (Molecular Probes/Life Technologies) for 10 min, washed with PBS, and finally coverslips were mounted in ProLong Gold antifade reagent (Life Technologies).

Individual RNAs were detected by fluorescence in situ hybridization using ViewRNA Cell Plus (Thermo Fisher Scientific, Cat#88-19000-99) and corresponding RNA probes (Thermo Fisher Scientific, Cat# VX-01), according to manufacturer’s instructions. Briefly, cells were seeded on 24-well plates with coverslips processed once they reach 80% confluency and fixed according to ViewRNA Cell Plus protocol. In some cases, for analysis of downregulated RNAs, cells were first fixed with 4% PFA for 1 h, followed by incubation with buffer A (PBS, containing 1 % TX100 and 4 μg/ml Proteinase K) for 10 min at RT. Cells were then washed 5 times with PBS, post-fixed with 4% PFA for an additional 1 h, washed with PBS and processed for hybridization.

For c-fos gene activation phorbol-12-myristate-13-acetate (524400, Sigma-Aldrich) in final concentration 1 ug/ml was applied for 2h to DLD-1^AID-NG-^Tpr cells prior to c-fos RNA visualization.

### RNA FISH microscopy and data analysis

poly(A)RNA fish image capturing and analysis was performed as described earlier^42^. Images were captured with a Zeiss Axiovert 200 M automated microscope using AxioVision software. A Zeiss 60X Plan-APOCHROMAT lens (1.4 NA) and Zeiss AxioCam MRm camera were used in imaging. Multiple 0.3 *μ*m z planes were captured and images were deconvolved with AutoQuant software. Imaris Surfaces tool was used for segmentation and signal analysis within the cytoplasm and nucleus. Images are representative of 18 images from two independent experiments. The graph provides the ratio of cytoplasmic to nucleus (C/N) poly(A) RNA with and without auxin treatment. Values represent means ± SD of at least 25 cells from biological triplicates. For poly(A) RNA accumulation at nuclear speckles cells were treated with 1 mM auxin for 8 h to degrade nucleoporins. After auxin treatment cells were fixed, stained with oligo^43^-Quasar 670 and anti-SON antibody. Images are representative of 9 images from biological triplicates.

Individual RNAs detected with ViewRNA Cell Plus were imaged and analyzed on Olympus IX71 inverted microscope utilizing an Olympus UPlanSApo 60x/1.4 oil objective as described in immunofluorescence staining section.

### Antibodies

The following antibodies were used for immunostaining and Western blot analysis: anti-mouse (A28175) and anti-rabbit (A11034) AlexaFluor-488 conjugated antibodies (Invitrogen); anti-mouse (A11004) and anti-rabbit (A11011) AlexaFluor-568 conjugated antibodies (Invitrogen); anti-mouse (A27042) AlexaFluor-680 conjugated antibodies (Invitrogen), anti-mouse (A-31553) AlexaFluor-405 conjugated antibodies. Specific primary antibodies against Nup50 (A301-782A), Nup153 (A301-789A), Tpr (A300-828A), GANP (A303-128A WB, A303-127A IF), NXF1 (A303-913A IF, A303-915A WB), Nup133 (A302-386A), Nup98 (sc-30112), HA (11867423001), FLAG M2 (F1804), Actin (13E5, 4970S), SON (GTX129778) were purchased from Bethyl Laboratories, Santa Cruz Biotechnology, Sigma-Aldrich, and GeneTex, respectively. Secondary HRP-conjugated anti-mouse and anti-rabbit antibodies for Western blot analysis were purchased from Sigma-Aldrich.

### Data access

The sequencing data of this study have been submitted to the NCBI Gene Expression Omnibus (GEO, http://www.ncbi.nlm.nih.gov/geo/) under accession ID GSE132363 (RNA-seq) and to the Sequence Read Archive (SRA) under BioProject ID: PRJNA548782 (ChIP Pol II Ser5P).

## Acknowledgements

VA, AA, SC, MD, CE, HNL and AS were supported by the Intramural Research Program of the *Eunice Kennedy Shriver* National Institute of Child Health and Human Development at the National Institutes of Health, USA (Intramural Project #Z01 HD008954). PB and BMAF were supported by NIH R01 GM113874-04 and R01AI125524-04. We thank NICHD Molecular Genomics Core for paired-end sequencing. We would like to thank Brian Brown, NIH Library Editing Service, for reviewing the manuscript.

## Author Contributions

VA, AA, MD developed hypothesis, conceived and designed the analysis. VA, MD wrote the paper. VA collected data and developed the project. SC, CE, VA, AS, PB carried out the experiments. HNL, JI performed bioinformatics analysis. VA, HNL, AS, SC, PB, BF, AA, MD edited manuscript.

## Completing Interests statement

None

## Figure Legends for Supplementary data

**Supplementary Figure 1.**
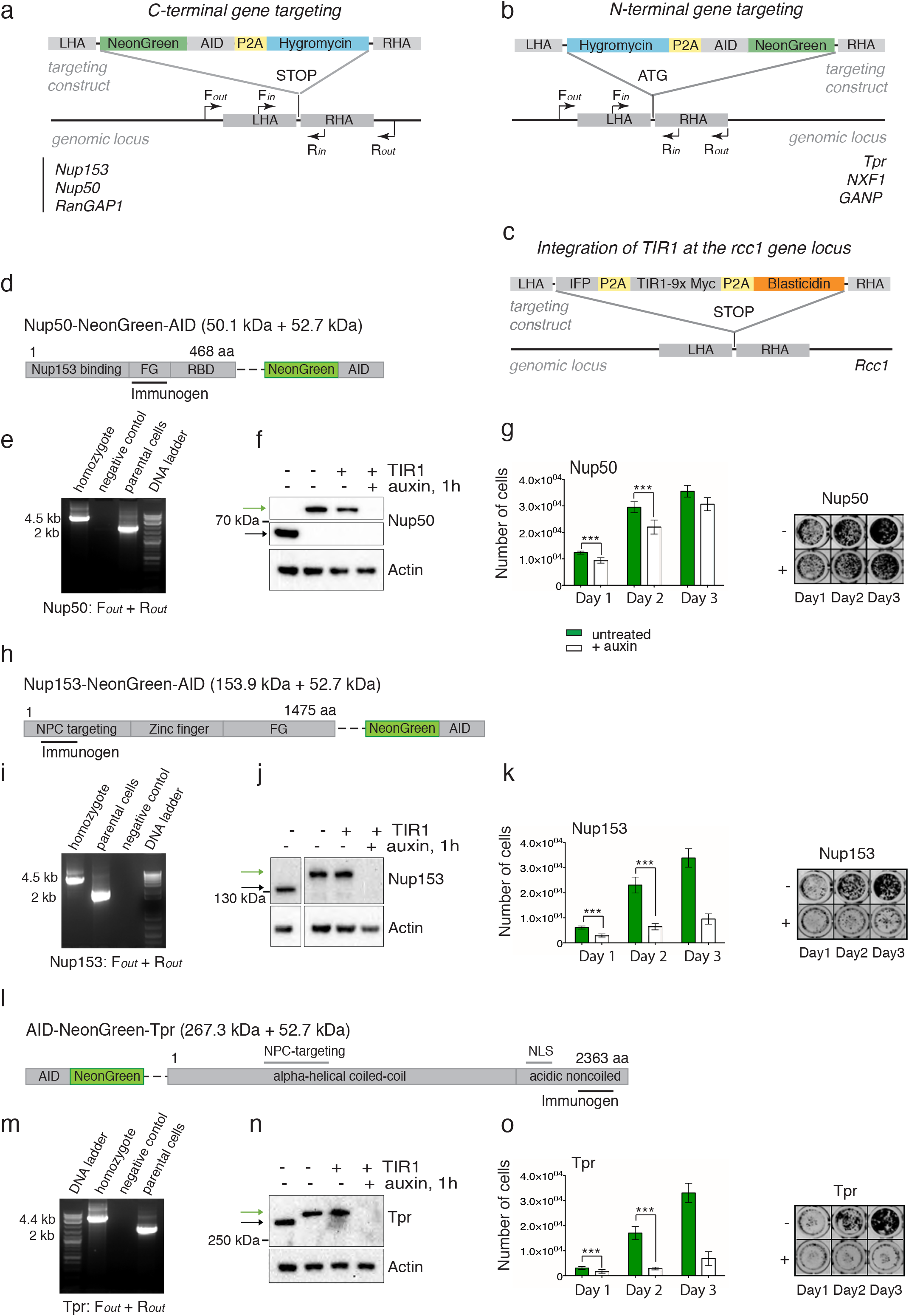
Degon targeting strategy. **a-b**, A scheme of endogenous targeting, position of homologous arms, and primer sets for genotyping of C-(a) and N-(b) terminally targeted genes. F_*in*_-R_*in*_ and F_*out*_-R_*out*_ primer sets anneal inside and outside of the homologous arms, respectively. We biallelically tagged Nup50, Nup153 and Tpr with an AID and a <u>N</u>eon<u>G</u>reen (NG) fluorescent protein, using CRISPR/Cas9 to modify their endogenous loci in DLD1 cells. **c**, A scheme of TIR1 genomic integration into the *rcc1* gene locus. **d, h, l**, Regions of Nup153, Nup50, and Tpr used for antibodies production (Bethyl Laboratories). **e, i, m**, Genomic PCR of homozygous clones demonstrating the integration of the NG-AID-P2A-Hygromycin sequence into genomic loci of *nup50, nup153,* and *tpr* genes. **f, j, n**, Degradation of Nup50 (**f**), Nup153 (**j**), and Tpr (**n**) after 1 h of auxin treatment. Black and green arrows indicate the molecular weight of unmodified and AID-NeonGreen (AID-NG) tagged nucleoporins. **g, k, o**, Growth rate of DLD-1 cells upon loss of Nup50, Nup153, and Tpr. Without auxin, AID-tagged cell lines were viable and did not show any obvious defects. Auxin addition led to rapid degradion of AID-tagged proteins and growth arrest of AID-Nup153 and AID-Tpr cells in the continuous presence of auxin. Error bars are SD. p-value *** < 0.0005 (unpaired t-test).

**Supplementary Figure 2.**
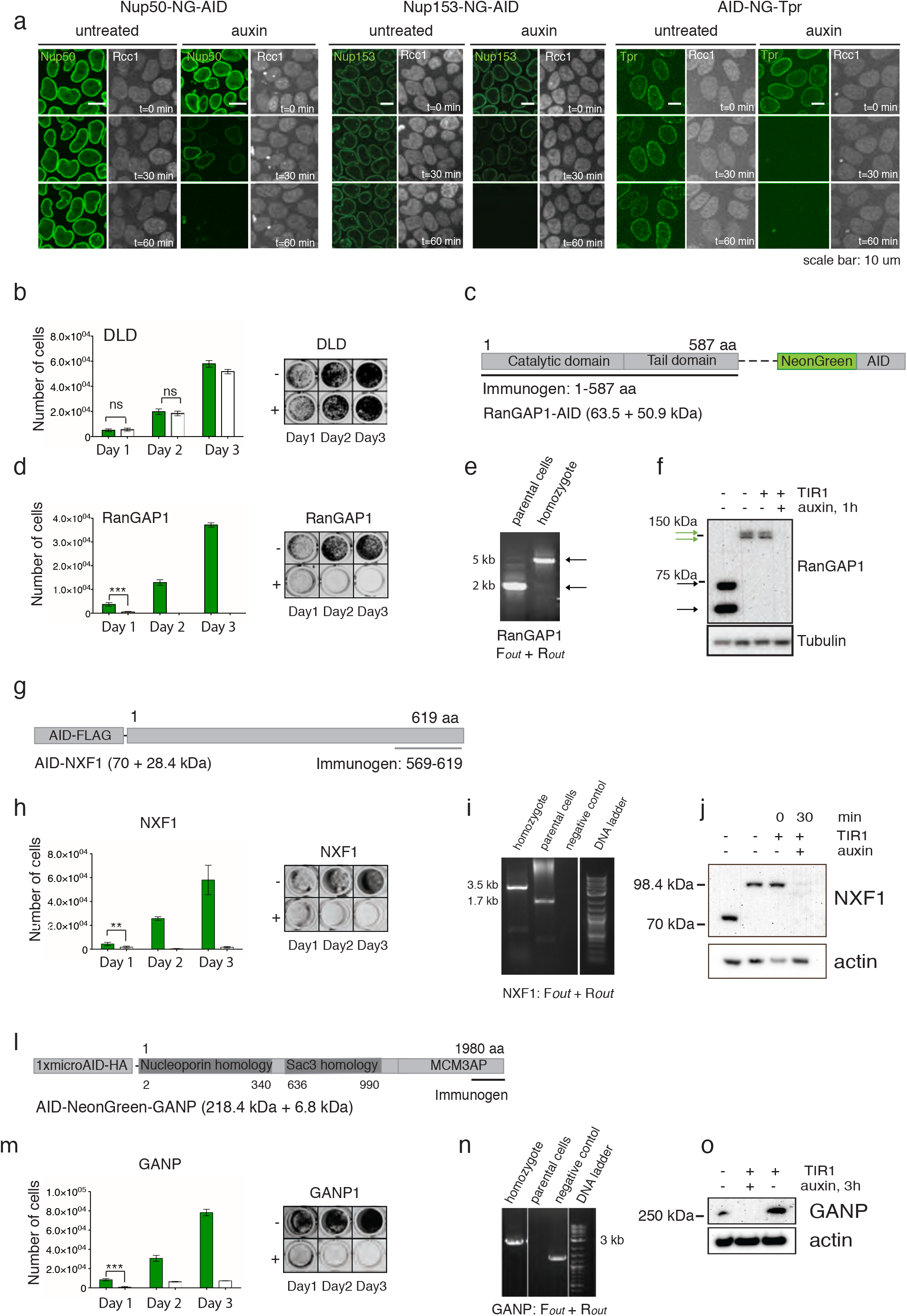
Degradation of BSK-NUPs. RanGAP1, NXF1, and GANP tagging strategy. **a**, Live imaging of Nup50, Nup153 and Tpr degradation. Chromatin is visualized by RCC1-IRFP fluorescent protein. Scale bar is 10 *μ*m. The green fluorescence intensity at the nuclear envelope corresponds to tagged nucleoporins in the absence or presence of auxin. The median degradation times for Nup50, Nup153 and Tpr were 60, 40, and 20 min, respectively. **b**, Growth rate of DLD-1 cells in the absence or presence of auxin. **c, g, l**, Protein regions of RanGAP1, NXF1, and GANP detected by respective antibodies. **d, h, m** Growth rates of AID-tagged RanGAP1, NXF1, or GANP1 cells upon auxin treatment for 1-3 days. Error bars are SD. p-value *** < 0.0005 (unpaired t-test). **e, i, n**, Genomic PCR demonstrating the integration of the AID degron sequence into the locus of *rangap1, nxf1*, or *ganp* genes. **f, j, o**, The molecular weight shift and degradation of endogenous RanGAP1, NXF1, and GANP were analyzed by Western blot upon auxin treatment. Black and green arrows indicate the molecular weight of endogenous and tagged proteins.

**Supplementary Figure 3.**
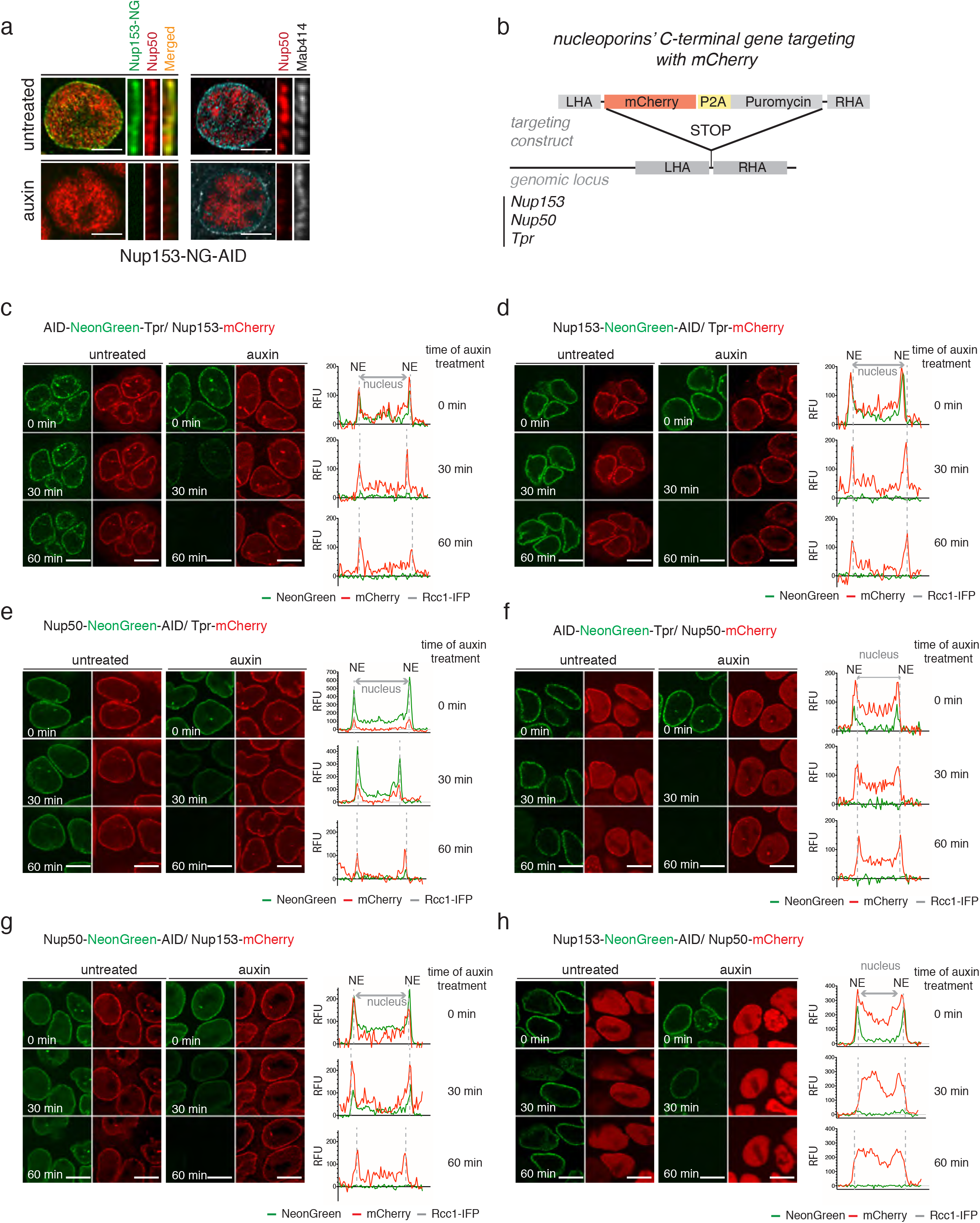
Stability of the assembled nuclear pores upon rapid loss of BSK-NUPs. **a,** Co-localization of Nup50 and Mab414-positive nucleoporins in the absence or presence of Nup153 (4 h of auxin treatment, scale bar is 5 *μ*m). **b**, A scheme of mCherry integration into genomic loci of *nup153, nup50,* and *tpr* genes. **c-h**, mCherry-tagged Nup153, Nup50 or Tpr were integrated into corresponding AID-BSK-NUPs cell lines and were used to follow Nup153-Tpr-Nup50 interdependency. The representative images of the mCherry signal at 30 and 60 min after auxin addition are shown. Scale bar is 10 *μ*m.

**Supplementary Figure 4.**
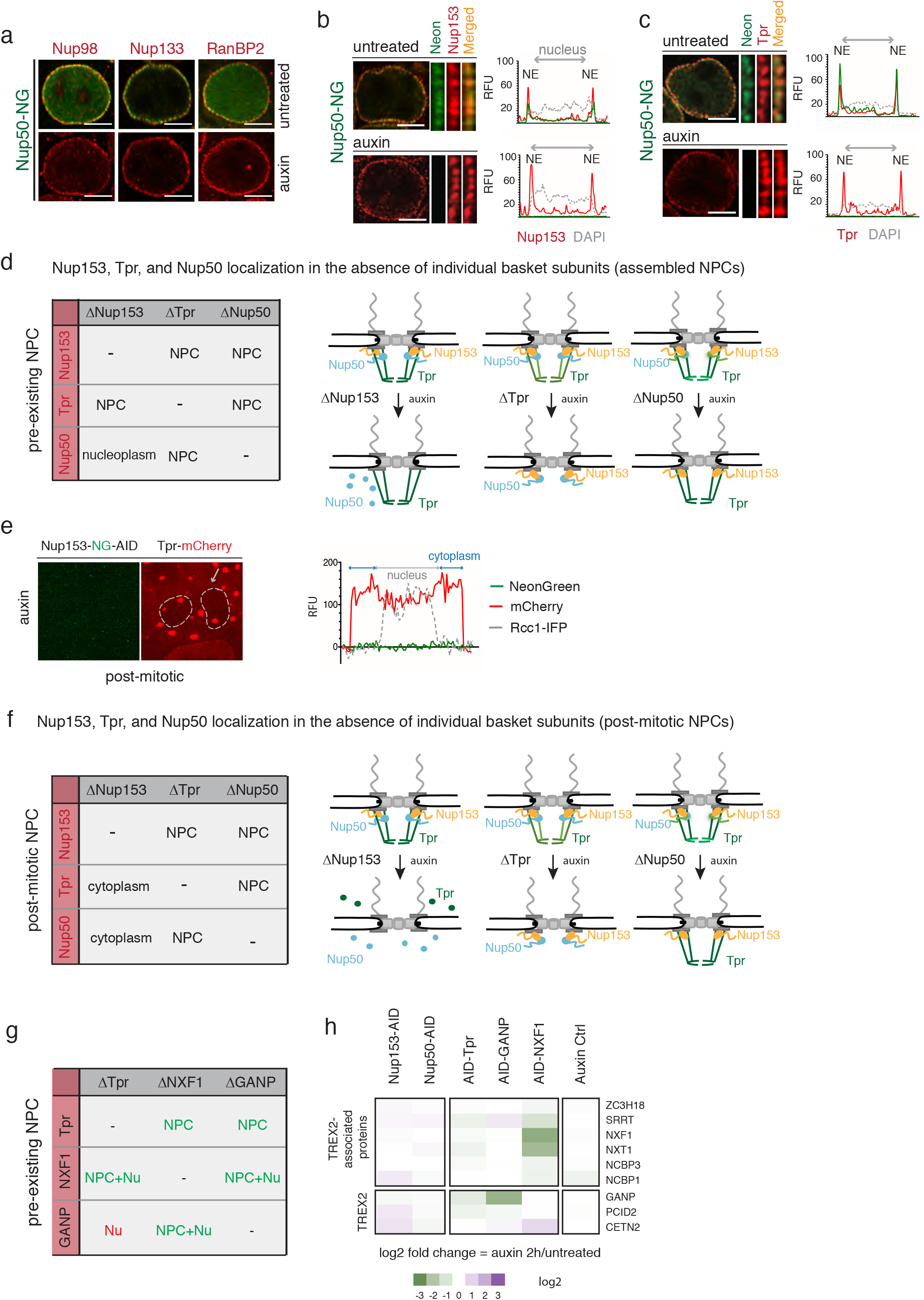
Supporting data regarding interdependence of BSK-NUPs. **a,** Localization of Nup98, Nup133, and RanBP2 nucleoporins on the NE in the absence or presence of Nup50 (4 h of auxin treatment). **b-c**, Localization of Nup153 (**b**) and Tpr (**c**) on the NE in the absence or presence of Nup50 (4 h of auxin treatment). Scale bar is 5 *μ*m. **d, f** Nup153-Nup50-Tpr interdependence in interphase (pre-existing NPC (d) and post-mitotic NPC (f) upon loss of Nup153, Tpr, and Nup50. **e**, Localization of mCherry-tagged Tpr in Nup153-depleted post-mitotic cells. Tpr does not mislocalize from the NE upon Nup153 loss in the preexisting NPC, but fails to be imported to the nucleus in post-mitotic Nup153-depleted cells. **g**, A summary table of Tpr, GANP, and NXF1 interdependence based on mass-spectrometry and immunofluorescence data. Red font indicates change after auxin treatment. **h**, A heat map of TMT-based mass-spectrometry of TREX2 complex and TREX-2 associated proteins abundance in NPC-enriched nuclear extracts upon loss of BSK-NUPs, GANP or NXF1.

**Supplementary Figure 5.**
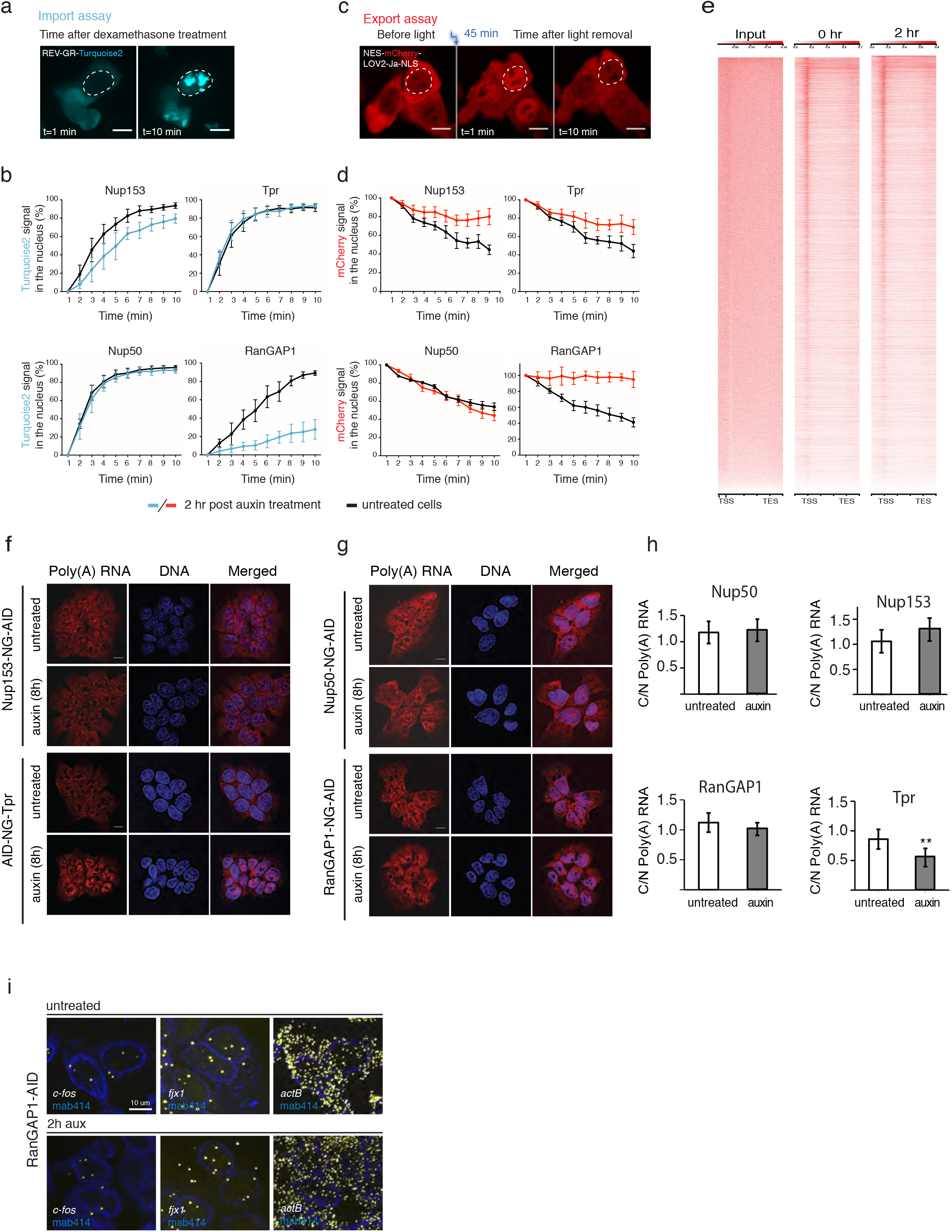
Impacts of BSK-NUP loss on nuclear-cytoplasmic transport and gene expression. **a, c**, Representative images of the import (**a**) export (**c**) assays. Scale bar, 10 *μ*m. In the import assay, a REV-GR-Turquiose2 substrate is localized in the cytoplasm. Upon treatment with dexamethasone, REV-GR-Turquiose2 becomes re-distributed to the nucleus. Import is measured by quantitation of Turquiose2 signal for 10 min immediately after dexamethasone addition. In the export assay, a mCherry-LOV2-BiLINuS2 substrate in loaded in the nucleus via blue light exposure. The mCherry-LOV2-BiLINuS2 shuttles back to the cytoplasm when blue light is switched off. Export is measured by quantitation of mCherry signal for 10 min immediately after removal of blue light. Import (**b**) and export (**d**) rates of model substrates in AID-tagged Nup50, Nup153, Tpr, and RanGAP1 cells in the presence (blue or red lines) or absence (black lines) of auxin. Error bars are MAD. **e**, A heat map of RNA Pol II Ser5P binding of AID-Tpr samples in the absence (0 hr) and presence (2hr) of auxin treatment. A region (± 2 kb) from transcription start site (TSS) and transcription end site (TES) is shown. Input samples did not show RNA Pol II Ser5P enrichment at the TSS or TES sites. **f**, Effect of BSK-NUPs and RanGAP1 depletion on the nuclear-cytoplasmic distribution of poly(A) RNA. Nup153, Tpr, Nup50, and RanGAP1 NG-AID targeted cells were analyzed using oligo(dT)-Quasar 670 probe. Cells were treated with 1 mM auxin for 8h. Representatives images from two independent experiments. Scale bar, 10 *μ*m. **h**, Poly(A) RNA export quantification in AID-tagged Nup50, Nup153, and Tpr cells in the presence or absence of auxin 8 h after auxin treatment. Error bars are SD. **< 0.005 for non-parametric t-test. **i**, *c-fos* and *fjx1* RNA localization in DLD1-RanGAP1^-NG-AID^ cells in the absence and presence of auxin. *c-fos* and *fjx1* RNA abundance is not changed upon RanGAP1 loss.

